# Variant Library Annotation Tool (VaLiAnT): an oligonucleotide library design and annotation tool for Saturation Genome Editing and other Deep Mutational Scanning experiments

**DOI:** 10.1101/2021.01.19.427318

**Authors:** Luca Barbon, Victoria Offord, Elizabeth J. Radford, Adam P. Butler, Sebastian S. Gerety, David J. Adams, Matthew E. Hurles, Hong Kee Tan, Andrew J. Waters

## Abstract

**Motivation:** Recent advances in CRISPR/Cas9 technology allow for the functional analysis of genetic variants at single nucleotide resolution whilst maintaining genomic context (Findlay et al., 2018). This approach, known as saturation genome editing (SGE), is a distinct type of deep mutational scanning (DMS) that systematically alters each position in a target region to explore its function. SGE experiments require the design and synthesis of oligonucleotide variant libraries which are introduced into the genome by homology-directed repair (HDR). This technology is broadly applicable to diverse research fields such as disease variant identification, drug development, structure-function studies, synthetic biology, evolutionary genetics and the study of host-pathogen interactions. Here we present the Variant Library Annotation Tool (VaLiAnT) which can be used to generate saturation mutagenesis oligonucleotide libraries from user-defined genomic coordinates and standardised input files. This software package is intentionally versatile to accommodate diverse operability, with species, genomic reference sequences and transcriptomic annotations specified by the user. Genomic ranges, directionality and frame information are considered to allow perturbations at both the nucleotide and amino acid level.

**Results:** Coordinates for a genomic range, that may include exonic and/or intronic sequence, are provided by the user in order to retrieve a corresponding oligonucleotide reference sequence. A user-specified range within this sequence is then subject to systematic, nucleotide and/or amino acid saturating mutator functions, with each discrete mutation returned to the user as a separate sequence, building up the final oligo library. If desired, variant accessions from genetic information repositories, such as ClinVar and gnomAD, that fall within the user-specified ranges, will also be incorporated into the library.

For SGE library generation, base reference sequences can be modified to include PAM (Protospacer Adjacent Motif) and protospacer ‘protection edits’ that prevent Cas9 from cutting incorporated oligonucleotide tracts. Mutator functions modify this protected reference sequence to generate variant sequences. Constant regions are designated for non-editing to allow specific adapter annealing for downstream cloning and amplification from the library pool.

A metadata file is generated, delineating annotation information for each variant sequence to aid computational analysis. In addition, a library file is generated, which contains unique sequences (any exact duplicate sequences are removed) ready for submission to commercial synthesis platforms. A VCF file listing all variants is also generated for analysis and quality control processes.

The VaLiAnT software package provides a novel means to systemically retrieve, mutate and annotate genomic sequences for oligonucleotide library generation. Specific features for SGE library generation can be employed, with other diverse applications possible.

**Availability and Implementation:** VaLiAnT is a command line tool written in Python. Source code, testing data, example library input and output files, and executables are available at https://github.com/cancerit/VaLiAnT. A user manual details step by step instructions for software use, available at https://github.com/cancerit/VaLiAnT/wiki. The software is freely available for non-commercial use (see Licence for more details, https://github.com/cancerit/VaLiAnT/blob/develop/LICENSE).

## 1 Introduction

The increase in human whole genome and whole exome datasets has led to the identification of millions of genetic variants. For example, the gnomAD consortium dataset contains sequence data for ∼200,000 individuals with close to 250 million high quality variants (Karczewski et al., 2020; Lek et al., 2016). Linking variants to disease states through Genome Wide Association Studies (GWAS) has furthered our understanding of gene function and the contribution of genetics to disease. For most variants, quantitative and qualitative differences relevant to disease association occur between individual carriers; this is likely due to haplotype, linkage disequilibrium, variable expressivity, incomplete penetrance and complex, pleiotropic, context-dependent phenotypes. This leads to a lack of statistical power, resulting in only a very small fraction of observed variants being conclusively interpreted (Starita et al., 2017; Weile & Roth, 2018). Furthermore, individuals with previously unobserved variants in known disease-risk loci face an uncertain genetic diagnosis if they carry alleles not obviously deleterious, with novel missense variants routinely categorized as ‘Variant of Uncertain Significance’(Cooper, 2015). Therefore, understanding the functional impact of variants is a key challenge of modern genomics research with implications for clinical management and our fundamental biological understanding of disease genes. This challenge has been met in part through the *in vitro* functional assessment of discrete, often clinically observed, variants through assays and cell culture experiments. However, the rate of discovery of new variants in disease loci calls for a proactive approach to functional assessment using high-throughput, Multiplexed Assays of Variant Effect (MAVEs) and Massively Parallel Reporter Assays (MPRAs) for coding and non-coding loci, respectively (Starita et al., 2017). A range of MAVE and MPRA technologies have been developed to produce variant effect maps, including yeast complementation (Hietpas, Jensen, & Bolon, 2011; Starr et al., 2020; Sun et al., 2016), cell display assays (Forsyth et al., 2013; Starr et al., 2020), FACs-based screens (Matreyek et al., 2018) including sort-Seq MPRA (Kinney, Murugan, Callan, & Cox, 2010), RNA-Seq MPRA (Melnikov et al., 2012) and CRISPR/Cas9-based saturation genome editing (SGE) in human cells (Findlay et al., 2018; Meitlis et al., 2020). A distinct benefit of SGE is that variants are assessed within the endogenous genomic context, allowing interrogation of complex mutational consequence, including splicing effects. This increases the relevance of these data for clinical interpretation.

In a typical cell culture SGE experiment, an sgRNA-Cas9 complex targeted to the genomic region to be edited induces a double stranded DNA break (DSB). The presence of co-transfected variant-harbouring repair templates lead to the incorporation of nucleotide changes at the locus through HDR. The cell population is then phenotyped, most commonly by assessing the fitness/growth of edited cells, and then analysed using deep amplicon sequencing.

Whilst the concept of SGE is straightforward, the design and production of variant libraries is not a trivial process. In order to facilitate the application of SGE to diverse applications we present a computational variant library generation tool (VaLiAnT) that allows straightforward *in silico* variant library production.

## 2 Methods and System

### 2.1 Overview

VaLiAnT is run from the command line. Input, output and broad processes are summarised in Fig. 1 and Supplementary Fig. 1. Reference files for genomic sequence (FASTA format), transcript features (GTF/GFF2 format) and custom variant accessions (VCF format) can be downloaded from repositories and stored locally. Multiple custom variant files can be used, with directories for VCF files listed in a custom variant manifest file (CSV format). Unique identifiers for the variant accessions will be extracted from the ‘ID’ column within the VCF files by default. Alternatively, unique identifiers can be parsed from user-specified INFO tags within the VCF files and included in metadata output.

**Fig 1.**
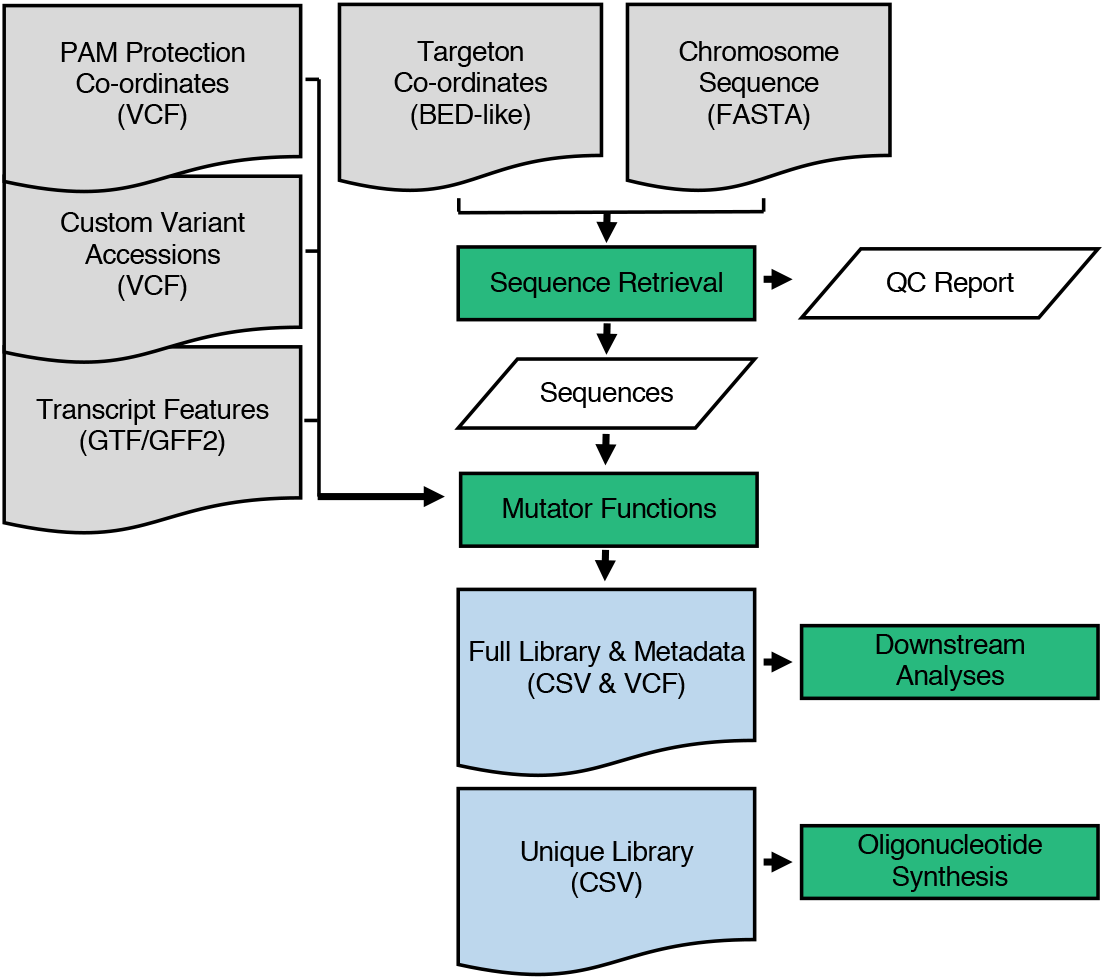
Information flow diagram for VaLiAnT: Input files are shown as grey document boxes with file types shown in brackets, processes are shown as green boxes and output files are shown as blue document boxes with file extensions in brackets. Process data is shown in slanted white boxes. Quality control (QC) report contains sequences in a comma-separated value (CSV) file and is execution-specific. All other output files are targeton-specific.

Depending on the nature of the experiment, the user may wish to target exonic, intronic or a mixture of both exonic and intronic sequences. To communicate this, we use the term ‘targeton’ to describe the region of the genome that will be targeted. For each targeton reference sequence, different mutator functions may be applied (Fig. 2). Basic mutation types do not consider coding sequence frame and are used for nucleotide-level saturation, actioned through ‘1del’, ‘2del0’, ‘2del1’ and ‘snv’ functions (Fig. 2a). CDS-specific mutation types consider the frame of the GTF/GFF2-specified transcript (Fig. 2b). Sitting between the nucleotide and protein saturation space, ‘snvre’ functions based on the amino acid-level mutational consequence (synonymous/missense/nonsense) of changes induced by ‘snv’ at the nucleotide level (‘snvre’ includes the ‘snv’ function) (Fig. 2b-i). The purpose of ‘snvre’ is two-fold. Firstly, to increase library representation of synonymous variants in SGE analyses. As synonymous changes have minimal functional impact on protein function they can act as base-line negative controls for read-count normalization. When ‘snv’ creates a synonymous change at a codon due to triplet degeneracy, ‘snvre’ creates all possible synonymous codons at the same triplet position (‘snv’ alone should create all 2-and 4-fold degenerate codons, i.e. when a 1 base-pair changes leads to all synonymous codons), expanding synonymous inclusion for 6-fold degenerate codons (arginine, leucine and serine) and stop codons (TGA>TAG and TAG>TGA).

**Fig 2.**
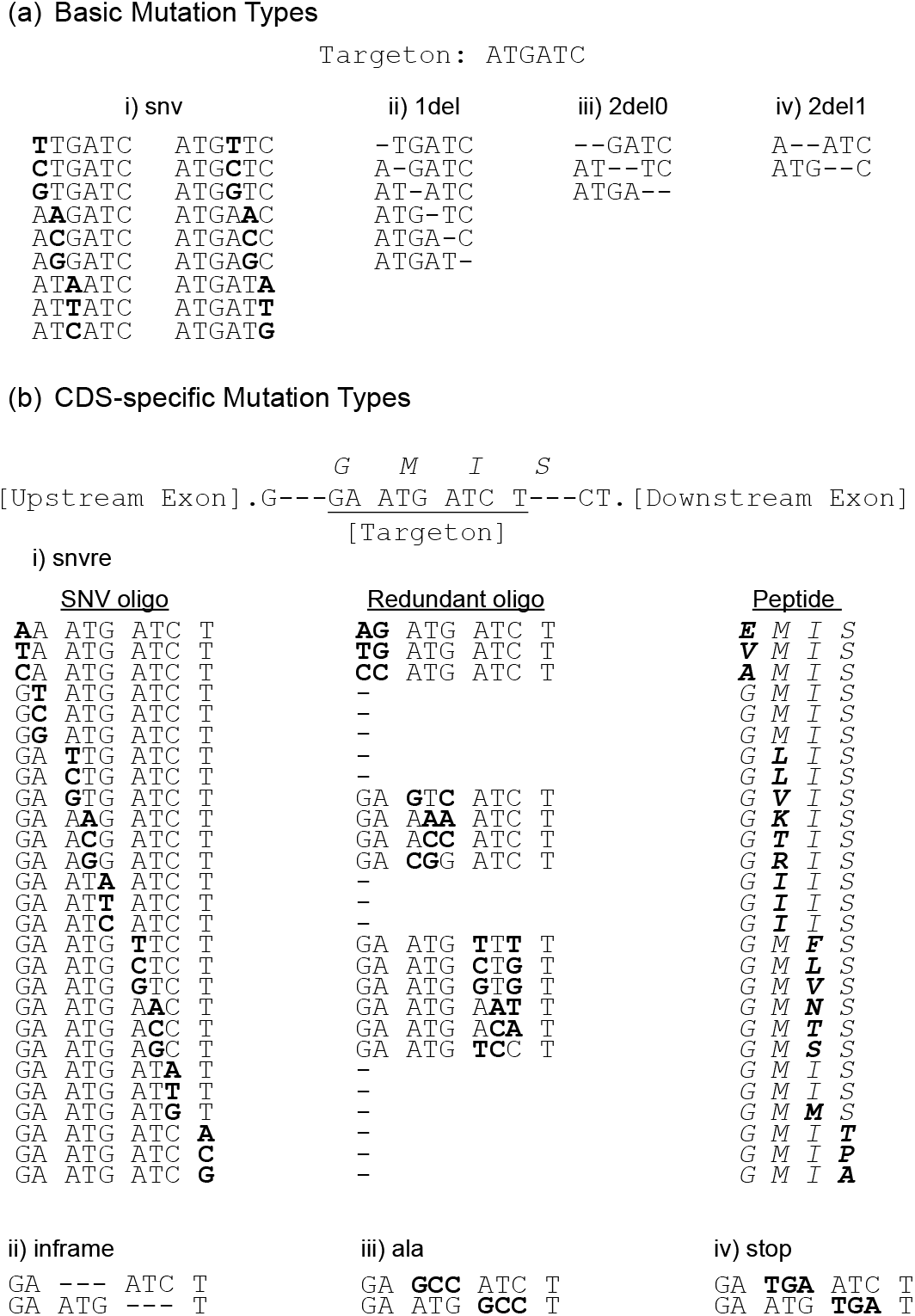
Mutator function descriptions: (**a**) Basic mutation actions that do not require CDS reading frame information. An example six base-pair targeton is shown for explanation purposes; a-i) ‘snv’ results in all possible nucleotide substitutions at each position in the targeton, a-ii) ‘1del’ deletes single nucleotides at each position, a-iii) ‘2del0’ deletes nucleotides in tandem starting at position 0, a-iv) ‘2del1’ deletes nucleotides in tandem starting at position 1. (**b**) Mutation actions that require CDS reading frame information. An example eight base-pair targeton is shown in which the first and last codon are split between upstream and downstream hypothetical exons. Amino-acids encoded by codons are displayed in capital italics above the DNA sequence; b-i) the ‘snvre’ mutation function runs the ‘snv’ function resulting in ‘SNV oligo’ output. CDS frame information is computed and where a missense change occurs as a result of ‘snv’ – shown in ‘Peptide’ in bold – a ‘Redundant oligo’ is generated. This redundant oligo encodes the same missense change as that generated by ‘snv’ at the peptide level, but with an alternative triplet sequence. The redundant triplet sequence chosen is the most frequent according to the codon reference table (or next most frequent if ‘snv’ generates the most frequent). Synonymous changes that result from ‘snv’ are included in ‘snvre’ outputs and are represented by ‘–’; b-ii) ‘inframe’ results in codon, triplet deletions for the longest in-frame coding sequence within the targeton; b-iii) ‘ala’ results in in-frame substitutions to alanine based on top-ranking alanine from the codon usage table, giving an alanine scan through coding sequence; b-iv) ‘stop’ results in in-frame substitutions to a stop codon based on the top-ranking stop from the codon usage table, giving systematic truncating mutations throughout the coding sequence.

Secondly, when ‘snv’ creates a mutation that leads to a missense change, the wild-type codon is exchanged for the next most frequent triplet code for the same missense change (based on a default or user defined frequency table), this allows for insight into concordant effects of missense change.

Protein level saturation is actioned through the ‘aa’ function. This function exchanges each wild-type codon for the most frequent triplet code of each other amino acid. While the default codon frequency table is human, VaLiAnT allows the user to provide an alternative codon frequency table. Nucleotide libraries that include both nucleotide and protein level full saturation of average exon sizes are highly complex. This may limit coverage by amplicon sequencing at desired levels (with current SGE protocols) (Sakharkar et al., 2002). Thus, VaLiAnT includes two functions as an alternative to the inclusion of full protein-level saturation sequences with nucleotide-level libraries, namely ‘ala’ and ‘stop’ (Fig. 2b-iii and iv respectively). These functions permit an in-frame alanine scan through coding sequences, and the replacement of codons with stop codons, respectively. In addition, we have included a function, ‘inframe’ that removes codons to create in-frame deletions (Fig.2 b-ii). Frame information for ‘aa’, ‘snvre’, ‘inframe’, ‘ala’ and ‘stop’ is derived from specific transcript GTF/GFF2 files provided by the user.

The user is likely to require different mutator functions for intron sequence and exon sequence. VaLiAnT has the option to split targetons into sub-regions to allow for different mutator functions to be enacted within different ranges of the targeton (Fig. 3a). This information is defined by the user in the form of a BED-like input ‘parameter’ file. Targeton ranges are defined by 1-based genomic coordinates using ‘ref_start’ and ‘ref_end’ fields, with reference chromosome and strand also defined as ‘ref_chr’ and ‘ref_strand’ respectively. Coordinates in fields ‘r2_start’ and ‘r2_end’ define a sub-region within the complete targeton range; typically, a complete or partial exon. A user-defined extension vector can then be applied to this range to define flanking sequence, upstream or downstream (relative to positive strand) of r2. Sequence defined by r2 and sequence within the extensible 5’ and extensible 3’ ranges can be mutated independently by mutator functions. An action vector split into three components, specified in the targeton input file, designates mutator functions to be enacted on the distinct sequence ranges in respective order (5’extension (r1), r2, 3’extension (r3). Any targeton sequence, defined in ‘ref_start’ to ‘ref_end’ but not in ‘r2_start’ to ‘r2_end’ or ‘ext_vector’ is included in oligonucleotide sequences as constant, unmodified genomic sequence (custom variants are still incorporated into constant regions). This is useful for defining flanking sequence for the annealing of downstream cloning adapters. In addition, generic adapters for universal amplification of all oligonucleotides in a synthesized pool can be specified at the command line and are appended after mutator function actions. Any generated sequences exceeding a user-defined length (default >300bp) are excluded from standard output files with a warning displayed to the user and separate metadata file returned.

**Fig 3.**
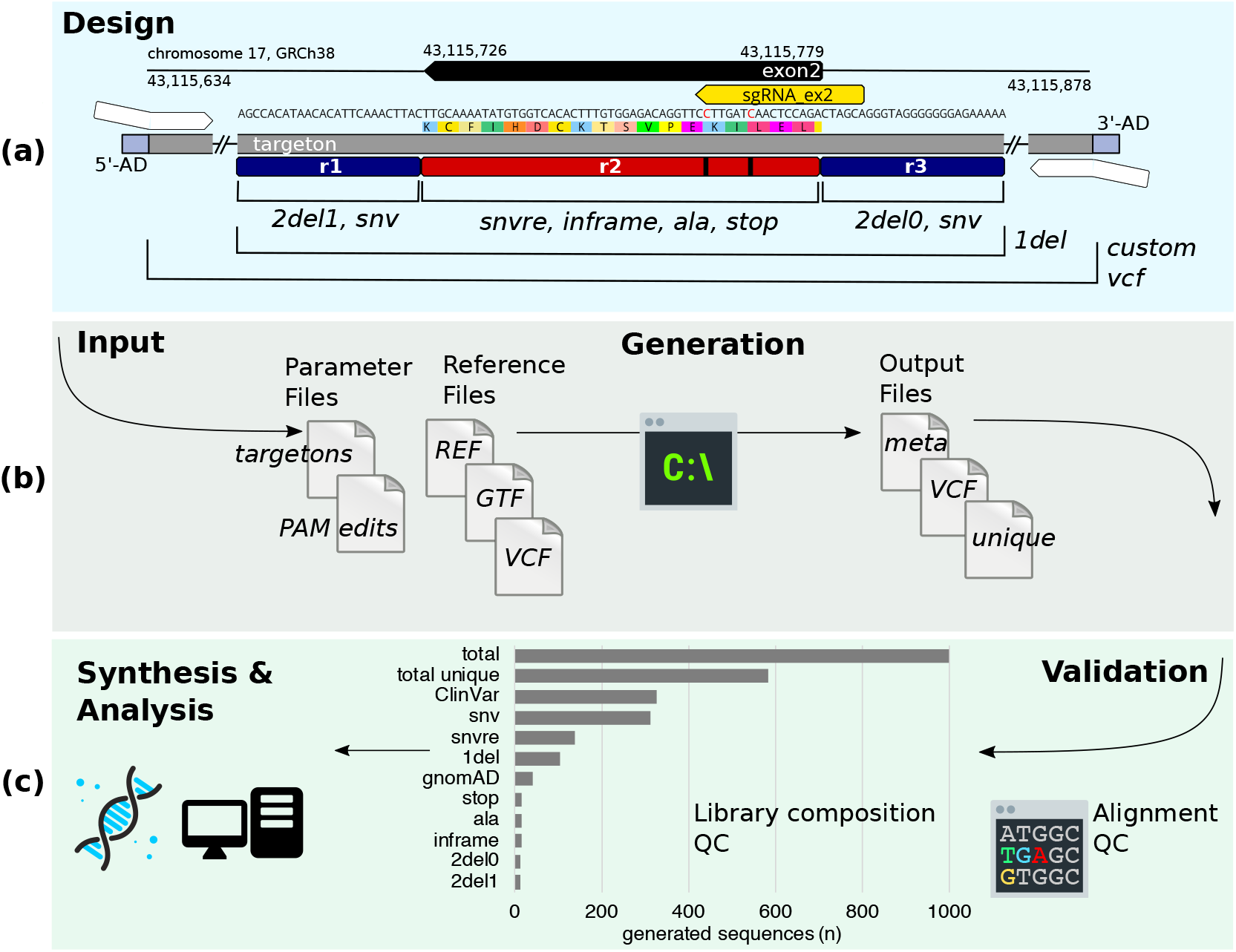
SGE library generation workflow for *BRCA1*: An example workflow for *BRCA1* exon 2 (*Homo sapiens*, GRCh38, ENSE00003510592) is shown with the corresponding input and output files used available at https://github.com/cancerit/VaLiAnT (**a**) Overview of the library design, with sequence information modified from Geneious Prime® (version 2019.04) visualization. Targeton, genomic region, chromosome and genome build are displayed, together with GRCh38 coordinates both for the complete targeton and for *BRCA1* exon 2. The selected sgRNA binding site is shown in yellow, where the directionality of the exon and sgRNA are negative strand and antisense respectively. Within the nucleotide sequence, red nucleotides are positions selected for PAM/protospacer-protection edits, beneath which the translated peptide sequence is shown by coloured rectangles. Below this, the targeton range is represented by a grey rectangle with condensed sequences represented by double slash. At either end of the targeton are the 5’-AD and 3’-AD (light blue rectangles) which represent appended P5 and P7 adapter sequences, enabling generic amplification of the generated library pool. Annealing sites for targeton-specific amplification and cloning adapters are shown as white arrows. Dark blue rectangles represent region 1 (r1) and region 3 (r3) and are 25bp extensions from region 2 (r2), a red rectangle with black lines to indicate the location of the PAM-protospacer protection edits. Regions in which mutator functions are actioned are described through annotated black demarcation lines. Variants ingested through custom VCF files are incorporated throughout the entire range of the targeton. As shown, r1 and r3 are modified with basic mutation type functions. To ensure deletion of dinucleotide splice-acceptor or splice-donor intronic sequences immediately flanking exon 2, sequential deletion through r1(25bp) is off-set by using 2del1 (as r1 length is odd), enabling deletion of exon-flanking tandem nucleotides at the distal (3’) end of r1. As dinucleotide deletion proceeds from the 5’ end, 2del0 is used for r3, ignoring the final distal nucleotide of r3 at the 3’ end. r2 is modified by CDS-specific mutator functions. (**b**) Schematic of the input and output files used for computation. The ‘targetons’ file contains targeton and r2 genomic ranges, r1 and r3 extension values and additional information including the required mutator functions for each region and sgRNA identifiers which correspond with the identifiers given in the ‘PAM edits’ VCF file. Reference files include ‘REF’ FASTA chromosome sequence, ‘GTF’ specific transcript annotation, and any custom variant files ‘VCF’. VaLiAnT is run from the command line to generate output files, including ‘meta’ full library metadata, ‘VCF’ of all variants generated and a ‘unique’ csv file for easy ordering of sequence synthesis. (**c**) Downstream processes. Interrogation of the output files using sequence alignment is used for library validation. A library composition graph is shown, delineating each mutator function output for the entire targeton range of *BRCA1* exon 2, including total sequences and total unique sequences. Downstream synthesis, experimentation and/or analysis processes are possible after validation.

To prevent Cas9 cleavage of variant sequences successfully incorporated at genomic loci, PAM and/or protospacer protection edits (synonymous/non-coding variants, refractory to Cas9-sgRNA cleavage) can be included in generated sequences. Protection edits are defined by the user in the form of a parameter VCF file, in which position, reference and alternative nucleotide required are defined, along with an sgRNA identifier. The protection variants are incorporated into each targeton tagged with the sgRNA identifier in the targeton input file, with all sequences generated for that targeton containing the protection edits. More than one protection variant per sgRNA and/or targeton is possible.

### 2.2 Implementation

The tool is implemented as a standalone executable Python package exposing a command line interface, requiring no Internet access. Tabular data is managed via the Pandas package and interoperation with bioinformatic file formats via the PySam (FASTA and VCF) and PyRanges (GTF/GFF2) packages. The tool is species and genome-build agnostic.

### 2.3 Candidate region selection and sequence retrieval

PAM/protospacer-protection edits are assigned via an sgRNA identifier to retrieved sequence defined by targeton range and used to create a modified reference sequence. The PAM/protospacer-protected reference sequence obtained is the basis for subsequent mutations (custom or generated). Discrete mutation events are applied per oligonucleotide so that each generated sequence contains specific PAM/protospacer protection edits and the desired variant composed of one or more base changes.

#### 2.3.1. Targeton configuration

Targeton genomic ranges and desired mutator functions are detailed in a BED-like tab-delimited file. Genomic ranges are expressed with the start position preceding the end position, regardless of the strand of the transcript. Each targeton can be divided into up to five regions (c1-r1-r2-r3-c2). Constant regions 1 and 2 (c1 and c2) are purposely not amenable to mutation through systemic mutator functions, but are still edited to include custom variants defined by VCF input. Regions1-3 (r1-3) can be changed by mutator functions–each independently of each other–by detailing corresponding mutator lists in the BED-like input file. Both r1 and r3 ranges are derived through extension from the defined r2 range, detailed by a numeric list in the input file. Optional PAM/protospacer-protection is configured via an sgRNA identifier list with corresponding variants retrieved from a dedicated VCF file.

#### 2.3.2. Reference annotation

In order to collect gene and transcript information and to apply CDS-specific mutator functions, appropriate transcript annotation must be provided via a GTF/GFF2 file; only CDS, UTR and stop features are taken into consideration. One transcript per gene is allowed to avoid ambiguities in matching target regions. No target region (r1, r2, r3) within a targeton can span both coding and non-coding sequences; although the complete targeton can be divided into coding and non-coding regions. Constant regions (c1 and c2) can span coding and non-coding regions if desired. UTR sequences are processed in the same way as intronic sequences. Targeton region reading frame is computed using user-specified transcript feature annotations. Retrieval of additional positions from the reference sequence is necessary to obtain the context of partial codons.

The frame of a sequence is obtained as follows, with *f* representing the frame as the number of bases missing from the codon at the 5’ end of a sequence, and *s* the start position:

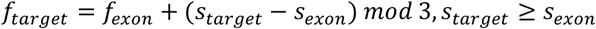

The number of additional positions 5’ and 3’ (0 ≤ *l*_*ext*_ ≤ 2), respectively, are obtained as follows, with *l* being the sequence length:

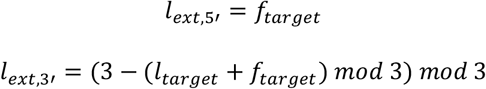

The extra bases required at either end of the target may come from the same or an adjacent exon (Supplementary Fig. 2).

#### 2.3.3. Reference sequence retrieval

Reference oligonucleotide sequences are retrieved from a local FASTA file. Unless a sequence starts at position one, the nucleotide preceding it is also retrieved; this is required to generate the REF and ALT fields of liminal variants in the output VCF file.

#### 2.3.4. PAM/protospacer-protection

The PAM/protospacer-protection step applies custom single-nucleotide variants that will be shared among all oligonucleotides for a targeton. The variant parameters are provided via a VCF file, each with an allocated SGRNA INFO tag, allowing variant grouping by sgRNA. sgRNA identifiers are assigned to targetons for specific inclusion.

### 2.4 Variant generation and output files

#### 2.4.1. Coding and non-coding variant generation

Variant incorporation can require different steps depending on whether the target region is a protein-coding sequence (CDS) or not. Basic mutation types, namely SNVs, single and tandem nucleotide deletions, can be introduced into either type of target; SNVs carry extra information when applied to CDS target regions. CDS-specific mutation types are applied to the longest in-frame CDS within the target or the shortest in-frame CDS containing the target (e.g. by considering distal nucleotides outside the target to complete partial, liminal codons, including bases from the preceding and/or following exon).

Variants that require the annotation of amino acid changes and mutation type (synonymous, missense, or nonsense) are generated from computed metadata tables.

The number of possible codons with a single SNV is limited, namely three per base per codon per strand (3 · 3 · 4^3^ · 2 = 1152). For the same codon on different strands, the nucleotide changes are the same but the amino acid changes, and therefore their classification (synonymous, missense, or nonsense), differ. If a mutator that requires SNV annotation is selected (‘snv’ or ‘snvre’), a partial metadata table, indexed by codon, is computed at start-up based on the codon table for each strand represented in the inputs. Similar considerations apply to the metadata of ‘snvre’ and codon substitutions. A partial table of all synonymous codon substitutions can also be generated from which a partial table of the top-ranking synonymous codon substitutions is subsequently derived.

#### 2.4.2. Custom variants

Custom variant VCF files are defined in a manifest list file. Simple variants are currently supported, including substitutions, insertions, deletions, and indels. In order to remain source agnostic, insertions and deletions are classified based on the POS, REF, and ALT fields. Start positions are corrected as appropriate before mapping variants to targetons. To preserve variant provenance, an alias is assigned to each VCF file. Optionally, an INFO tag mapping to an identifier (e.g. ALLELEID for ClinVar variants) can also be assigned. By default, the VCF ID field is used as the variant identifier. Variants that start and end within the targeton are applied, generating one oligonucleotide sequence each.

#### 2.4.3. Oligonucleotide sequence output

Either three or four targeton-specific output files and one sequence quality control (QC) file (reporting retrieved wild-type sequence for the specified ranges) are generated per execution. The file names report chromosome, coordinates, strand, and the sgRNA IDs associated with the targeton (Supplementary Table 2 and Supplementary Table 3).

Variant oligonucleotide sequences are generated by replacing target regions in PAM/protospacer reference sequence with mutated sequence. The final oligonucleotide sequences are then assembled from: invariant region sequences (optional), adaptor sequences (optional) and target region mutated sequences. All variants are applied to the positive strand sequence. For negative strand transcripts, reverse-complemented final targeton sequence (excluding any adapter sequence) can be generated (optional) for ease of interpretation or for experimental requirement. The directionality of sequence for single-stranded synthesis will not affect downstream SGE experiments if PCR amplification (resulting in double-stranded DNA HDR template) forms part of library processing. In the case of oligonucleotides exceeding a specified, configurable length (e.g. due to a long insertion variant), metadata is segregated and sequences are not included in the unique list (a fourth output file containing excluded sequence information is generated).

#### 2.4.4. Output files and formats

Supplementary Table 2 lists output files and their main use. It should be noted that the output metadata format does not follow VCF convention in reporting positions, reference, and alternative sequences to favour streamlining of downstream processing. These departures only apply to deletions and insertions:

- deletions start at the first deleted position, the reference sequence starts at the same position, no alternative sequence is reported;
- insertions start at the position following the last unaffected nucleotide, the alternative sequence starts at the same position, and have no reference sequence.

Reference and alternative sequences for variants starting at position one similarly do not include the nucleotide that immediately follows.

## 3 Results

### *BRCA1* Nucleotide and Peptide Saturation Libraries

As an example of workflow and output we have generated oligonucleotide libraries suitable for performing SGE experiments on the exons encoding the RING domain of the tumour-suppressor gene *BRCA1* (*Breast cancer type 1 susceptibility protein*). From the transcript NM_007294.4 (ENST00000357654.9, Gencode release v34), exons 2, 3, 4 and 5 are targeted. A summary of the process for exon 2 of *BRCA1* is shown in Fig. 3. All input and output files are available at https://github.com/cancerit/VaLiAnT. Input coordinates consist of GRCh38 genomic ranges within *BRCA1* (ENSG00000012048.23, Ensembl release GRCh38.p12) for four targetons spanning each of exons 2, 3, 4 and 5 and PAM/protospacer protection edits at one sgRNA binding site per targeton. Input coordinates for exon 2 are shown in Fig. 3a and input criteria for exon 2 shown in Table 1. sgRNAs and PAM/protospacer protection edits were selected based on design parameters outlined in Supplementary Table 1. Targetons 2, 3 and 4 are 245 bp in length, covering the complete CDS of the exon with 25 bp extensible regions (r1 and r3) to target flanking intronic sequence. Targeton 5 is 251 bp in length with extensible regions of 20 bp (r1) and 41 bp (r3) at 5’ and 3’ of the CDS respectively. Nucleotide and amino acid saturation libraries have been created independently (brca1_nuc and brca1_pep respectively), with all mutator functions except ‘aa’ represented in the nucleotide library and ‘aa’ and ‘inframe’ in the amino acid library. Custom variants using ClinVar (release 2020-11-07, (Landrum et al., 2018)) and gnomAD (v3.0, (Karczewski et al., 2020)) VCF files for *BRCA1* have been incorporated in brca1_nuc (VCF files were filtered for genomic coordinates of interest to reduce file size). Exon 2 library complexity is summarized in Table 2 with a summary of the complexity of each library provided in Supplementary Table 3. P5 and P7 sequences were selected as adapter sequences to be added 5’ and 3’ of each oligonucleotide generated. Sequences that exceed 300 bp were omitted from unique sequences files using the ‘max-length’ option. As *BRCA1* is transcribed from the negative strand, the ‘revcomp-minus-strand’ option was used to generate reverse complement sequence. This does not affect correct sequence generation, it is a preference for validating generated sequences intuitively.

**Table 1.** Summary of input parameters for *BRCA1* exon 2 SGE library generation: Summary of values included in targeton parameter input file for exon 2. Values correspond to Fig. 3a schematic.

| input parameter | value |
| --- | --- |
| *chromosome* | 17 |
| *strand* | - |
| *targeton range* | 43115634 : 43115878 |
| *r2 range* | 43115726 : 43115779 |
| *r1 length* | 25 |
| *r3 length* | 25 |
| *sgRNA identifier* | sgRNA_ex2 |
| *mutators for r1* | 2del1, snv, 1del |
| *mutators for r2* | snvre, inframe, ala, stop, 1del |
| *mutators for r3* | 2del0, snv, 1del |

**Table 2.** Summary of output sequence categories for *BRCA1* exon 2 library generation: values for each attribute are shown, specific to regions 1-3 (r1-3) and constant regions (unedited, except for custom variants) and the summed values comprising the entire targeton. One custom variant in r2 results in an oligo longer than 300bp and is excluded from the final library, total unique oligos – the number representing library complexity for SGE experiments – is shown in bold.

| mutator functions |  | r1 | r2 | r3 | constant regions | complete targeton |
| --- | --- | --- | --- | --- | --- | --- |
|  | *sequence length (bp)* | 25 | 54 | 25 | 141 | 245 |
|  | *2del1* | 12 | – | – | – | 12 |
|  | *2del0* | – | – | 12 | – | 12 |
|  | *1del* | 25 | 54 | 25 | – | 104 |
|  | *inframe* | – | 17 | – | – | 17 |
|  | *snv* | 75 | 162 | 75 | – | 312 |
|  | *snvre* | – | 140 | – | – | 140 |
|  | *ala* | – | 17 | – | – | 17 |
|  | *stop* | – | 17 | – | – | 17 |
|  | *gnomAD* | 2 | 8 | 10 | 22 | 42 |
|  | *ClinVar* | 74 | 176 | 76 | 1 | 327 |
|  | *total* | 188 | 591 | 198 | 23 | 1000 |
|  | *excluded (>300bp)* | 0 | 1 | 0 | 0 | 1 |
|  | *total unique* | 109 | 351 | 101 | 22 | 583 |

## 4 Discussion

As the technology to write DNA catches up with our ability to read DNA, having the ability to computationally generate multiplexed variant libraries from wild-type sequence is a critical starting point for many studies. We present VaLiAnT, a software package that is immediately useful for those who wish to perform SGE experiments. Furthermore, VaLiAnT is agnostic with respect to species, genome build, transcript and custom variant source. The definition of targeton sequence range and the delineation of sub-regions within the targeton is versatile and can be modulated to suit experimental needs. Optional, user-defined PAM/Protospacer protection edits required to prevent Cas9 (or alternative Cas protein) cleavage of incorporated HDR template at target loci can be incorporated into all targeton output sequences.

We have created a range of functions to introduce changes within oligonucleotide sequences. Mutator functions systemically edit sequence to allow investigation in both nucleotide and protein space, which overlap to some degree for most functional assays. For example, clinical interpretation of a single nucleotide variant that leads to a missense change inherently involves drawing conclusions about the effect of a change in the protein sequence. In terms of questions that can be addressed through deep mutational scanning, the relative importance of nucleotide and protein level saturation can be roughly attributed. Clinical interpretation and prediction of variant effect, including splice sites, will mostly be informed by saturating at the nucleotide level; fundamental biological investigation of structure-function relationships and drug target development will likely benefit most from saturation at the protein level; and studies in pathogen and evolutionary biology will benefit from interpretation of both nucleotide and protein level saturation.

There are several reported methods to create nucleotide variant libraries using either error-prone PCR or forms of chemical synthesis. While error-prone PCR is comparatively cheap, variants cannot be systemically generated to the same degree as with oligonucleotide-array, chemical synthesis platforms. *In vitro* enzymatic generation of oligonucleotides using the template-less DNA polymerase terminal deoxynucleotidyl transferase is being pursued by several biotechnology companies with some encouraging developments, and might prove to be an effective alternative to chemical synthesis in the future (Liberante & Ellis, 2021). At the time of writing chemical synthesis at the scale of ≤300nt is possible on a large scale. Undoubtedly, as synthesis platform technology advances, an increase in oligonucleotide synthesis length (and reduction in error rate) will be become possible. We have provided an optional filter to remove sequences above a user-defined length, which is configurable to accommodate for future increases in synthesis capability.

VaLiAnT has been designed to aid in the design and generation of oligonucleotide libraries for SGE experiments. However, VaLiAnT has a wide range of possible uses including, but not limited to: mutational consequence annotation of coding-sequence variation, conversion of VCF annotation files to oligonucleotide sequences and the generation of libraries for exogenous or orthogonal assays (as opposed to genome editing), that use cDNA cassettes and/or transcriptional readouts (Jones et al., 2020; Ursu et al., 2020). The reverse-complement option in VaLiAnT permits the generation of variant sequences for strand-specific applications. For example, SGE using single-stranded oligo donor (ssODN) has been reported to streamline experimental procedures, however strand-specific differences in HDR-rate have been observed with ssODN template and orientation should be chosen depending on editing context (Kan, Ruis, Takasugi, & Hendrickson, 2017; Meitlis et al., 2020). Therefore, VaLiAnT could be used to generate strand-specific libraries for ssODN-based SGE experiments.

Prime-editing based saturation mutagenesis is another strand-specific application in which VaLiAnT could be used. Reverse-transcriptase (RT) templates, which contain the desired variants and are strand-specific, form part of prime-editing guide RNAs (pegRNAs) (Anzalone et al., 2019). VaLiAnT could be used to produce saturating edits in RT template and the adapter option used to append scaffold and guide RNA spacer sequence identified with pegFinder, producing pegRNA oligonucleotide sequences for synthesis and subsequent saturation mutagenesis (Chow, Chen, Shen, & Chen, 2020).

Saturation genome editing (SGE) provides rich information for functional analysis of genetic variation as variants are assayed in their endogenous genomic environment. As SGE and other DMS studies with next-generation sequencing readouts increase in number, it is anticipated that tissue/cell-specific studies, will provide further context and biological relevance in addition to genomic environment. Tissue/cell specific analysis of variants and their effects on protein structure and protein-partner complexes is also an area that DMS will greatly inform. Inferring protein structure from DMS data is possible as is imputation of probable variant effects from a mutational spectrum (Dunham, Beltrao, Molecular, & Campus, 2020; Schmiedel & Lehner, 2019). In addition, as predicative models and machine learning increase in utility for characterization of the properties of mutational change and prediction of the likely effect of mutational change, particularly in relation to protein-protein interactions, training datasets rich in biological context will be important (Senior et al., 2019). Therefore, being able to use specific genomic reference sequence and distinct transcript information in the design of variant libraries – which VaLiAnT allows – will become necessary.

We envision that, as open source software, VaLiAnT will be further modified by the community. For example, by the addition of new mutator actions to expand the repertoire of available functions useful for different forms of experimentation, annotation or analysis. There is also scope for VaLiAnT to be combined with other software, such as upstream heuristic functions to select appropriate input information, such as; targeton ranges, sgRNAs and PAM/protospacer protection edits. In addition, VaLiAnT may be combined with downstream analysis software for bioinformatic analysis of deep mutational scanning data.

VaLiAnT provides a novel way to systemically produce *in silico* variant libraries and corresponding annotations and metadata. We believe the software package will become an essential component of the deep mutational scanning toolkit, significantly contributing to the study of genetic perturbations at scale.

## Acknowledgements

We would like to thank Leopold Parts, Sofia Obolenski and Mark Thomas for their help in testing the software and for commenting on the manuscript.

---

**Supplementary Fig. 1.**
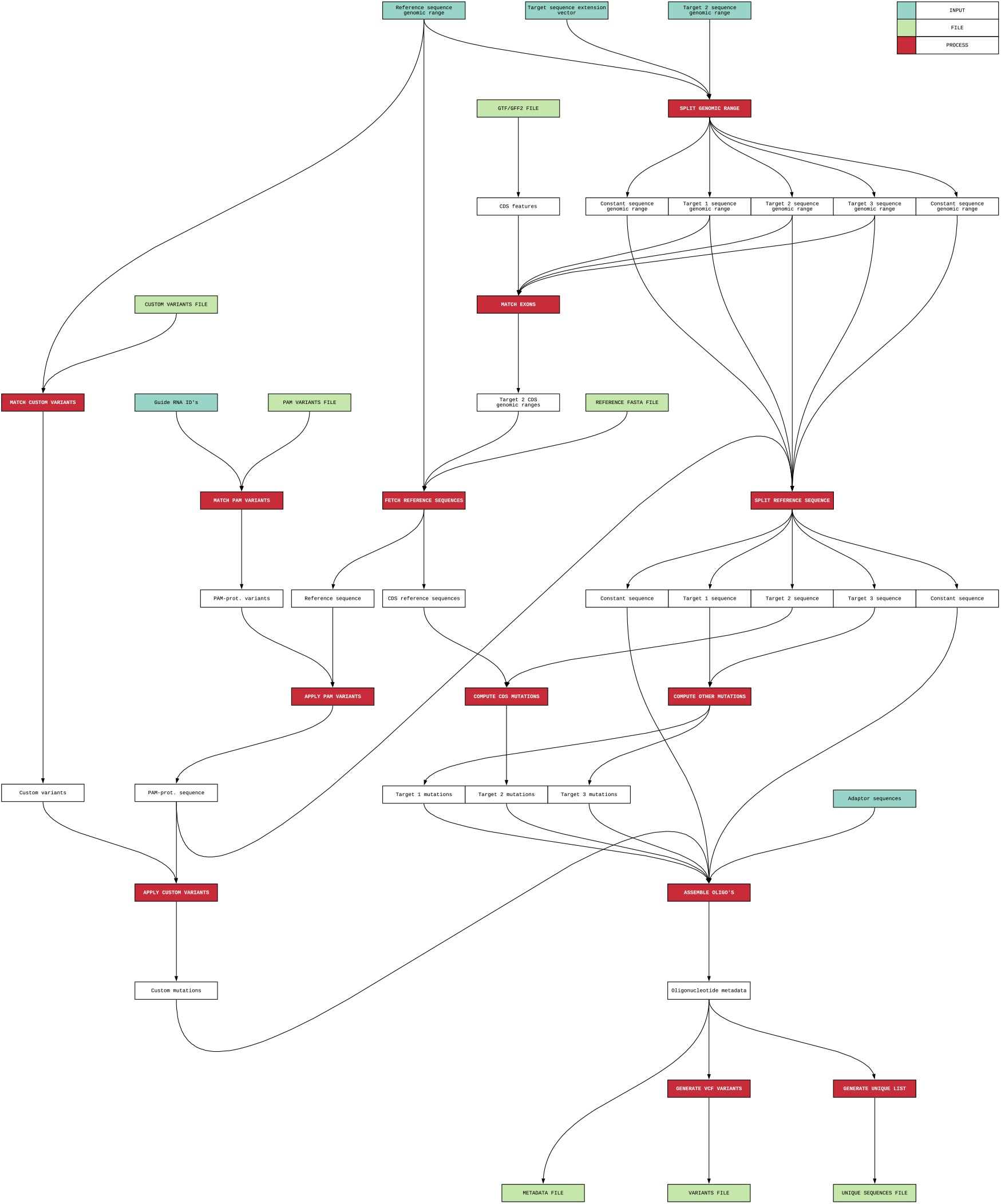
Flow of information through VaLiAnt: inputs (‘INPUT’) are represented by dark green rectangles, files (‘FILE’) by light green boxes and processes (‘PROCESS’) by red rectangles.

**Supplementary Fig. 2.**
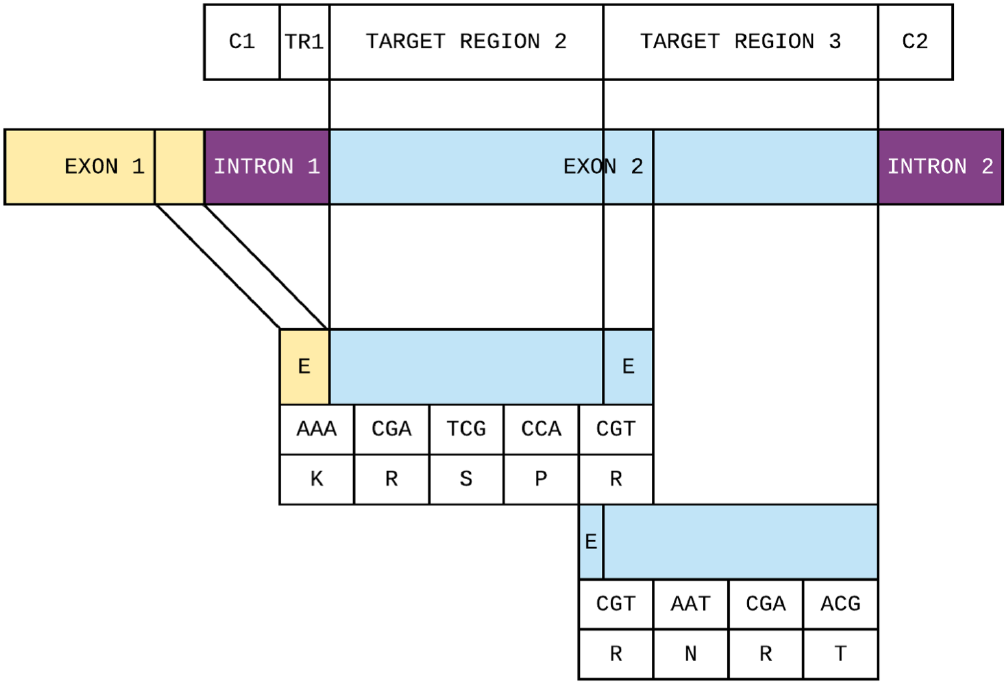
Reading frame calculation logic: an example targeton with the first constant (C1) and target (TR1) regions on the first intron of a transcript, the second and third target regions on the second exon, and the second constant region (C2) on the second intron. The 5’ CDS extension for TR2 is retrieved from the preceding exon, the 3’ CDS extension from the same. TR3 requires no CDS extension at 3’ because the second exon ends in-frame.

**Supplementary Table 1:** sgRNA site and PAM/protospacer protection nucleotide selection criteria: Summary of attribute considerations and reasons driving selection of sgRNAs for *BRCA1* exon 2-5 SGE library design and nucleotide selection for PAM/protospacer protection edits within the sgRNA binding site sequence.

| sgRNA selection criteria |  | PAM/protospacer protection criteria |  |
| --- | --- | --- | --- |
| *attribute* | *reason* | *attribute* | *reason* |
| sgRNA binding site at appropriate locus. | Fits design parameters. | Only mutate 3<sup>rd</sup> base of a codon at PAM site or in protospacer. | Codon redundancy allows for synonymous edits. |
| sgRNA binding site within targeton CDS (if targeting exon). | Frameshifting INDELs created by NHEJ will deplete (if CDS mutation is deleterious) increasing relative representation of HDR edits in screen. | Always change pyrimidines to pyrimidines (C/T) and purines to purines (A/G). | Missense changes caused by SNV saturation at PAM/protospacer protected codons will replicate missense changes achieved as if codon were wild-type (with some exceptions*). |
| Synonymous edit possible at PAM and/or protospacer. | Prevents Cas9 from targeting HDR library template, which depletes representation of edited loci. | Wherever possible select a pyrimidine base to change. | No exception to above missense representation. |
| No 0 (exact), 1 or 2 mismatch to CDS off-target. | Avoids off-target effects and preserves on-target activity. | Wherever possible change terminal G of PAM over more protospacer proximal G. | Avoids NAG PAM sites which spCas9 still cuts with limited efficacy. |
| No 4 (or more) consecutive thymine residues (TTTT). | Prevents premature transcriptional termination of sgRNAs expressed from RNA polymerase III promoters (such as U6::3). | Antisense sgRNAs are preferred (CCN PAM rather than NGG). | An increased number of pyrimidine changes are possible. |
| Avoid sgRNAs that have G/GG adjacent to PAM NGG (leads to two consecutive GG at PAM after protection edits). | Despite PAM editing to protect from Cas9, cutting still occurs if NGG exists adjacent to wild-type PAM. | No protection edits in the unique codons, ATG[M] and TGG[W] or ATA[I]. | Purine protection edits at the 3<sup>rd</sup> position of these codons do not lead to synonymous changes. |

*\*in addition to a purine change at 3<sup>rd</sup> base, if: 1<sup>st</sup> base=T or 2<sup>nd</sup> base=G, then W/STOP missing in SNV SGE library; 1<sup>st</sup> base=A or 2<sup>nd</sup> base=T, then M/I missing in SNV SGE library; 1<sup>st</sup>+2<sup>nd</sup> base=TT or 1<sup>st</sup>+2<sup>nd</sup> base=AG, then both W/STOP and M/I will be missing in SNV SGE library.*

**Supplementary Table 2:** Summary of output file nomenclature and use: Details are shown for the five possible output files for a VaLiAnT processed targeton. For each output file, ‘filename’ shows the final descriptive extension appended to the targeton file names (see https://github.com/cancerit/VaLiAnT for the *BRCA1* exon 2 example output files). Broad and non-exclusive ‘main uses’ are noted. All files except the ‘ref_sequence.csv’ sequence retrieval QC file (which is produced per execution and contains information on all inputted targetons) are generated per targeton. Exclusion metadata files are only produced when a generated sequence exceeds 300bp (default) or user-specified maximum length.

| filename | description | main use | targeton-specific | execution-specific |
| --- | --- | --- | --- | --- |
| _meta.csv | Metadata and sequence annotation file | Bioinformatic analyses | ✓ | – |
| _meta_excluded.csv | Metadata file for sequences above specified length | QC of sequence exclusion | ✓ | – |
| _unique.csv | Filtered file containing only unique sequences and oligo name | Synthesis submission | ✓ | – |
| _.vcf | Variant Call Format of generated and custom variants | Bioinformatic analyses | ✓ | – |
| ref_sequences.csv | Retreived sequences for targetons and sub-regions | QC of sequence retrieval | – | ✓ |

**Supplementary Table 3:** Summary of library complexity for generated *BRCA1* libraries: ‘library’ name corresponds to exon and saturation space, ‘generated name’ contains targeton-specific generated nomenclature to describe library parameters, ‘saturation space’ is either nucleotide or amino acid-level, nucleotide mutator functions are as described in Fig. 3a for each of exon 2, 3, 4 and 5. Amino acid-level contains only ‘aa’ and ‘inframe’ mutator functions directed at the same targeton ranges and regions as the nucleotide libraries. ‘targeton length’ is the length in base-pairs of the entire targeton, ‘saturation length’ is the length in base-pairs of the sequentially mutated regions within the targeton (that is r1, r2, r3 combined), ‘total sequences’ is the number of oligonucleotide sequences produced (including multiples of identical sequence), ‘excluded sequences’ is the number of sequences exceeding the maximum length (default of 300bp), ‘unique sequences’ is the number of unique oligonucleotides generated.

| library | generated name | saturation<br>space | targeton<br>length | saturation<br>length | total<br>sequences | excluded<br>sequences | unique<br>sequences |
| --- | --- | --- | --- | --- | --- | --- | --- |
| brca1_nuc_ex2 | *chr17_43115634_43115878_minus_sgRNA_ex2* | nucleotide | 245 | 104 | 1000 | 1 | 583 |
| brca1_nuc_ex3 | *chr17_43106355_43106599_minus_sgRNA_ex3* | nucleotide | 245 | 128 | 1209 | 1 | 740 |
| brca1_nuc_ex4 | *chr17_43104794_43105038_minus_sgRNA_ex4* | nucleotide | 245 | 139 | 1292 | 0 | 825 |
| brca1_nuc_ex5 | *chr17_43104080_43104330_minus_sgRNA_ex5* | nucleotide | 251 | 201 | 1439 | 1 | 1185 |
| brca1_pep_ex2 | *chr17_43115634_43115878_minus_sgRNA_ex2* | amino acid | 245 | 54 | 340 | 0 | 340 |
| brca1_pep_ex3 | *chr17_43106355_43106599_minus_sgRNA_ex3* | amino acid | 245 | 78 | 500 | 0 | 500 |
| brca1_pep_ex4 | *chr17_43104794_43105038_minus_sgRNA_ex4* | amino acid | 245 | 89 | 580 | 0 | 580 |
| brca1_pep_ex5 | *chr17_43104080_43104330_minus_sgRNA_ex5* | amino acid | 251 | 140 | 920 | 0 | 918 |

